# Annotation of Phage Genomes with Multiple Genetic Codes

**DOI:** 10.1101/2022.06.29.495998

**Authors:** Aaron Pfennig, Alexandre Lomsadze, Mark Borodovsky

## Abstract

Some of recently discovered in human gut microbiome highly divergent crAssphages were reported to use multiple genetic codes. Opal or amber stop codon reassignments were present in parts of the genomes, while the standard genetic code was used in the remaining genome sections. Essentially, the phage genomes were divided into distinct blocks where one or another code was used. We have developed a tool, Mgcod, that identifies blocks with specific genetic codes and annotates protein-coding regions. We used Mgcod to scan a large set of human metagenomic contigs. As a result, we identified hundreds of contigs of viral origin with the standard genetic code used in some parts while genetic codes with opal or amber stop codon reassignments were used in others. Many of these contigs originated from known crAssphages. Further investigation revealed that while the genes in one genomic block could be translated by a distinct genetic code, translation of genes by either of the two genetic codes genes in an adjacent block would produce proteins with little difference from each other. The dual-coded genes were enriched with early-stage phage genes, while a single code was used for the late-stage genes. The code-block structure expands the phage’s ability to infect bacteria whose genomes employ the standard genetic code. The new tool provides means for accurate annotation of unusual genomes of these phages.

## Introduction

The discovery of the genetic code (Matthei et al. 1962; Nirenberg 2004) was announced as a revelation of a fundamental truth universal across all domains of life (Crick 1968). Still, after a relatively short time, a presence of deviations from the ‘standard’ genetic code assignments was discovered in many organisms (Barrell, Bankier, and Drouin 1979; Yamao et al. 1985; Caron and Meyer 1985; Syozo Osawa and Jukes 1989; S. Osawa and Jukes 1995).

The advent of massive NGS-based deciphering of genomes of free-living organisms and viruses brought in a necessity for computational tools able to identify codon reassignments along with genome annotation. Several such tools, designed to identify a genome-wide species-specific genetic code, use multiple alignments of homologous nucleotide and amino-acid sequences (Abascal, Zardoya, and Posada 2006; Dutilh et al. 2011; Mühlhausen and Kollmar 2014; Noutahi et al. 2017; Shulgina and Eddy 2021). However, alignment-based methods are less effective for deeply branching species; therefore, an *ab initio* approach is also needed.

Another complication with variations of genetic codes was revealed in studies of crAssphages discovered in human gut microbiomes (Dutilh et al. 2014; Yutin et al. 2018; Guerin et al. 2018; Edwards et al. 2019; Koonin and Yutin 2020; Benler et al. 2021). It was shown that in some crAssphages genera, the *opal* (TGA) or *amber* (TAG) stop codons are not used in the same way in the whole genome. The genomes could be divided into distinct regions (blocks) that differed in terms of the type of employed genetic code (Ivanova et al. 2014; Guerin et al. 2018; Yutin et al. 2021; Borges et al. 2021). Obviously, a genome with multiple genetic codes is hard to correctly decode by the conventional genome annotation tools (Ivanova et al. 2014; Borges et al. 2021).

Here, we describe a new tool – Mgcod – that combines recognition of genetic code, parsing of phage genomes into blocks with differences in genetic codes, and gene prediction. We focus on genetic codes defined by stop codon reassignments, as sense-to-sense codon reassignments do not affect the performance of the genome annotation algorithms. Mgcod segments phage genomes with multiple genetic codes into blocks with block-specific genetic codes. This approach enables accurate annotation of coding regions in phage genomes with multiple genetic codes, such as crAss-like phages.

We used Mgcod for screening a set of human (gut) metagenomes. We pointed out 739 viral contigs with genetic codes characterized by reassignments of the *opal* and *amber* stop codon. Out of these 739 viral contigs, 222 appeared to be fragments of genomes of known crAss-like phages, while 517 were likely to belong to previously unknown crAss-like phages. All these contigs had blocks with block-specific genetic codes. Consistent with previous findings, we found that the switch of genetic code closely follows a switch in the protein-encoding strand (PES). We also observed that in the genomes with genomic block structure one genomic block encoded early-stage phage genes, while the other block was enriched for late-stage phage genes (Ivanova et al. 2014; Yutin et al. 2021; Borges et al. 2021). In contrast to previous studies that predicted distinct genetic codes in each block, we found that while a distinct genetic code (e.g., code A) was used in one block, the use of two different codes for translating genes in another block (code A and code B) would yield nearly identical proteins. Thus, genes in the second block appeared to be dual-coded. Notably, the dual-coded block was usually enriched for early-stage phage genes, while the other block was enriched for late-stage genes often encoded with a single genetic code.

## Materials

### Data sets

#### Genomes from databases

To assess the ability of Mgcod to correctly identify different genetic codes, we used 100 bacterial genomes with *opal* stop codon reassignment present in the RefSeq database as of September 2019 (Table S1) (O’Leary et al. 2016). In 99 genomes, TGA was reassigned to tryptophan (genetic code 4), and in one genome, TGA was reassigned to glycine (genetic code 25). All genomes with *opal* stop codon reassignment had a size less than 2Mb and a GC-content (GC%) between 20% and 35%. We also used 95 bacterial genomes with standard genetic code (genetic code 11) whose size and GC% are similar to that of genomes with *opal* stop codon reassignment (Table S2).

To ensure that the predictions of Mgcod are not biased by the low GC% of the training genomes, we also used 100 bacterial genomes with standard genetic code, sizes greater than 2Mb, and GC% greater than 35% (Table S3). We used these genomes to simulate a set with the stop codon reassignment for evaluation of the performance of Mgcod (see below).

In what follows we refer to *opal* stop codon reassignment as genetic code 4 (representing both TGA reassigned to tryptophan and TGA reassigned to glycine), to *amber* stop codon, TAG, reassignment to glutamine as genetic code 15 (reusing the discontinued NCBI nomenclature), and to *ochre* stop codon reassignment to tyrosine as genetic code 101 (introducing an ID designation for TAA reassignment in otherwise standard genetic code).

#### Simulation of genomes with genetic codes 4, 15, 101

No bacteria have genomes with *amber* stop codon or *ochre* stop codon reassignments. Furthermore, it was observed that bacterial genomes with *opal* stop codon reassignment have lengths larger than 2Mb, and GC% >35%. For the sake of completeness of the modeling, we generated artificial sequences with all types of reassignments, genome length, and GC%. For instance, to simulate an artificial genome with genetic code 15, we took one of the representative genomes with standard genetic code and predicted genes with MetaGeneMark using the standard genetic code model (Zhu et al.2010). Then, predicted TAG stop codons were replaced with either TGA or TAA. Next, 80% of the in-frame codons coding for glutamine, i.e., CAA and CAG, were substituted with TAG. This approach was supported by the observation that in real genomes with genetic code 4, TGA makes up 80% to 90% of the codons coding for tryptophan.

#### Simulation of contigs with blocks featuring multiple genetic codes

To assess the accuracy of Mgcod, we used simulated genomic constructs. To simulate genomes with multiple genetic codes, we used 100 genomes of bacteria with *opal* stop codon reassignment (genetic codes 4 and 25) as well as 95 bacterial genomes that employ the standard genetic code and are similar in size and GC%. A genome with either the standard genetic code or *opal* stop codon reassignment was randomly selected, and an arbitrary segment with a length between 25 kb and 175 kb was spliced out. Then, a genome with the other genetic code was randomly selected, and a random segment was cut. The length of the second segment was chosen such that the total length of the two segments would be up to 200 kb (which is similar to the size of crAss-like phages with multiple genetic codes (Guerin et al. 2018; Yutin et al. 2018)). If the randomly chosen excision point fell into a protein-coding region, it would be moved to the center of the adjacent intergenic region. To simulate the PES bias previously observed in phages with multiple genetic codes (Ivanova et al. 2014; Yutin et al. 2021; Borges et al. 2021), we *post hoc* updated the predicted strand of coding regions such that the switch of genetic code coincides with a switch of PES. Protein-coding regions were predicted with MGM. Due to automatic adjustment of the model parameters to local sequence composition, this algorithm worked without additional training on these synthetic fragments having different codon usage patterns in their parts (Zhu et al. 2010).

#### Metagenomes

We used Mgcod to search 6474 metagenomes deposited in GenBank in November 2019 for contigs with stop codon reassignments. Being interested in phage genomes, we selected 361,992 contigs with a length between 80 kb and 300 kb. Previously described phages with multiple genetic codes had a length between ∼95 kb and ∼180 kb (Dutilh et al. 2014; Guerin et al. 2018; Yutin et al. 2018).

## Methods

### Filtering of metagenomic contigs with multiple genetic codes

To create a high confidence set of metagenomic contigs with multiple genetic codes, a series of filtering steps were applied. First, we filtered out contigs that did not originate from human (gut) metagenomes and whose viral nature was not confirmed by VirFinder (*p-value* ≤ 0.05; Ren et al. 2017). The reasoning for this filtering step is that multiple genetic codes have only been observed in certain phages, and those phages were found to be highly abundant in human metagenomes (Dutilh et al. 2014; Guerin et al. 2018; Yutin et al. 2018). Also, we excluded contigs that were predicted with more than two blocks and in which the shorter block made up less than 20% of the contig. Finally, we identified contigs that represent putatively complete phage genomes based on circularity, i.e., an exact match of 50 – 200 bp at the ends. This set of putatively complete phage genomes constituted the high confidence set of phage genomes with multiple genetic codes.

### Generating evidence for predicted stop codon reassignments in phage contigs

To validate stop codon reassignments predicted by Mgcod, we attempted to build clusters of homologous nucleotide sequences of coding regions predicted in phage genomes with multiple genetic codes. We used UCLUST v11 in *cluster_fast* mode with an identity threshold of 0.8 to find clusters of homologous sequences (Edgar 2010). Nucleotide sequences within each cluster were aligned using Clustal Omega (v1.2.4) (Sievers and Higgins 2018). Then, the number of times a putatively reassigned stop codon was aligned to a particular amino acid or to other stop codons was counted, excluding gap symbols. If predicted stop codon reassignments are correct, one would expect that in MSAs of homologous sequences, the reassigned stop codon predominantly aligns to sense codons coding for the same amino acid or to the same type of stop codon.

### Functional annotation, tRNA prediction, and pairwise average nucleotide identities

Protein-coding regions in putatively complete phage genomes with multiple genetic codes were extracted and translated according to their predicted genetic code. Subsequently, the amino acid sequences were split based on their genetic code and clustered with an identity threshold of 0.9 using UCLUST (v11) in *cluster_fast* mode (Edgar 2010). The cluster centroids were then functionally annotated with hmmscan (v3.3) searches against the PVOG database (Grazziotin, Koonin, and Kristensen 2017; Mistry et al. 2013), and only the top-hit per query was retained after applying e-value cut-offs of 0.0001 and 0.01 for the full query and the protein domain, respectively. Furthermore, putatively complete phage genomes with multiple genetic codes were searched for suppressor tRNAs using tRNAscan-SE (v2.0.5), applying the bacterial model (-B) and a bitscore cut-off of 35 (Chan et al. 2021). Suppressor tRNAs predicted as “pseudo” were excluded from consideration. Pairwise average nucleotide identities (ANIs) between putatively complete and incomplete phage genomes with multiple genetic codes were determined using fastANI (v1.1) (Jain et al. 2018). Only the best match per query among matches with ANIs ≥ 0.8 was reported.

### The Mgcod Algorithm

Mgcod is a fully automated pipeline that recognizes stop codon reassignments, detects multiple genetic codes in phage genomes, parses the genome into blocks with different codes, and annotates protein-coding regions. In this section, after a brief overview, we describe individual parts of the algorithm in greater detail.

We start with a run of MetaGeneMark (Zhu et al. 2010) with four different models of protein-coding regions: i) the standard genetic code (genetic code 11), ii) a model with the *opal* stop codon reassigned to tryptophan (genetic code 4), iii) a model with the *amber* stop codon reassigned to glutamine (genetic code 15), and iv) a model with the *ochre* stop codon reassigned to tyrosine (genetic code 101). We chose MetaGeneMark as a tool for delivering accurate gene prediction in metagenomic sequences (Zhu et al. 2010). To predict a uniform genetic code, the code model that yields the highest coding potential for the entire genomic sequence (characterized by log-odds score) is chosen, and the corresponding gene annotation is recorded. To predict multiple genetic codes, we apply a sliding window approach, and for each window, we determine the model of protein-coding regions that has the highest coding potential. If one model has the highest coding potential in all windows, the corresponding genetic code is assigned to the entire contig. Otherwise, the genome is segmented into blocks with distinct genetic codes based on window labels indicating the highest-scoring genetic code model in the window. The genetic codes specific to each block are used to generate gene annotations (see Algorithm S1 in Supplementary Materials).

The final segmentation of a contig into blocks is done based on the assigned window labels and PES information. To avoid fragmentation of a genome into many short blocks, a minimum of *n* consecutive windows with the same label are required to form a block (by default, *n=3*). Therefore, sets of windows with the same label interrupted by fewer than *n* windows with a different label are merged. The merging procedure includes four steps: First, the longest sequence of consecutive windows with the same label is taken as a seed. The corresponding label is set as a reference. Second, this seed is extended in both directions until *n* consecutive mismatches are observed. Third, labels of windows in the extended segment are updated according to the reference label. Fourth, excluding windows with already updated labels, steps one to three are iteratively repeated until the labels of all windows have been updated. Then, the center of the intergenic region separating two blocks is defined as the boundary. Finally, the predictions of block boundaries are refined by searching for a change of PES in the proximity of the predicted block boundary. If a change of PES is observed in either one of the two windows bordering the predicted block, the block boundary is moved to the center of the intergenic region, separating the genes between which the PES switch occurs. The use of the PES information for segmenting phage genomes is justified by the observation that in phages with multiple genetic codes, the switch of genetic code coincides with a switch of PES (Ivanova et al. 2014; Yutin et al. 2021; Borges et al. 2021).

Mgcod’s ability to identify the correct model of protein-coding regions in a window depends on the window size. The accuracy of the identification declines with decreasing window size as less data – fewer genes – are available to render a decision. However, the resolution in terms of detecting multiple genetic codes decreases with increasing window size. We found that a window size of 5000 bp provides accurate identification of the genetic code while delivering sufficient accuracy in detecting genetic code variations along the genome (Table S4).

If multiple genetic codes are predicted, gene annotations are compiled from predictions of the two models that most frequently have the highest coding potential in a window. The predictions of these two models are compared, and for each gene, a decision of whether to include it in the final gene set is made based on the following rules: i) If one model predicts a long gene that spans multiple short genes predicted by the other model in the same PES, and the coding potential of the longer gene is greater than the aggregated coding potential of the short genes, the long gene is included in the final gene set, and the short predictions are dropped. ii) If the predictions of both models are in the same PES and share the stop coordinates or share the start coordinates while the stop coordinates differ by less than *t* nucleotides (by default, *t=30*), the predictions are considered equivalent. In these cases, the translation with either genetic code yields nearly identical proteins, i.e., the gene is dual-coded. For this reason, predictions of both models are included in the final gene set as isoforms. We consider predictions equivalent when the stop coordinates match because sometimes the models predict slightly different start coordinates due to marginally different emission probabilities. Similarly, if a gene does not have in-frame stop codon occurrences and if a model of an alternative genetic code predicts only a slightly longer gene compared to the other model, we consider the predictions equivalent. Such alternative genes are likely to be equivalent from the functional point of view because it is unlikely that a small extension of the coding region would have a significant impact on the protein function. iii) If the predictions of both models are identical, the gene is dual-coded, but only one prediction is included in the final gene set. iv) A gene uniquely predicted by one model is included in the final gene set.

When predicting multiple genetic codes, the sliding window approach is beneficial for the following reason. The algorithm was observed to be more robust when the segmentation into blocks was done based on the window labels rather than the gene labels indicating the genetic code with the highest coding potential. Segmenting a contig based on gene labels turned out to create too many blocks.

Mgcod takes a multi FASTA file as an input and generates gene annotations for each contig in GTF/GFF2 format. Additionally, the output comprises a tab-separated file indicating the number of reassigned stop codons per gene and the corresponding coordinates, a tab-separated file listing the segmentation of each contig into blocks with different genetic codes, and a visualization of the contig segmentation. Mgcod is available at https://github.com/gatech-genemark/Mgcod. We also provide a script for running the analysis of multiple contigs on a multiprocessor machine, thus scaling up the analysis of metagenomes.

## Results

### Detection of multiple genetic codes by Mgcod

To assess how well Mgcod detects multiple genetic codes, we simulated 1000 genomes with blocks of standard genetic code (code 11) and genetic code 4 (see Materials and Methods). Mgcod predicted correct alternation of genetic codes in 982 (98.2%) genomes. In the remaining 18 genomes (1.8%), Mgcod predicted either a single genetic code or too many changes of genetic code along the genome. Furthermore, in the 982 genomes in which Mgcod correctly predicted multiple genetic codes, it also accurately predicted the genomic region where the code switch occurs. The average shift in the predicted switch position was “-166” bp with a standard deviation of 5174 bp, suggesting that Mgcod predicted the switch close to the correct position of the switch with a standard deviation of one sliding window (5000 bp). Mgcod predicted the switch with a deviation of less than 5000 bp in 870 (87.0%) genomes (Table 1).

**Table 1:** Performance of Mgcod on 1000 simulated genomes with blocks of standard genetic code and genetic code 4. The number of genomes is shown in parenthesis.

|  | Overall (1000) | 11 → 4 (493) | 4 → 11 (507) |
| --- | --- | --- | --- |
| **Correct sequence of genetic codes identified** | 98.2% (982) | 98.4% (485) | 98.0% (497) |
| **Mean error in bp** | -166 | 47 | -374 |
| **Standard deviation in bp** | 5174 | 4634 | 5644 |
| **Mean error ≤ 5000 bp** | 87.0% (870) | 88.6% (437) | 85.4% (433) |

Mgcod’s accuracy in locating the switch of genetic code might differ depending on whether the genetic code 4 block follows the standard genetic code block or vice versa (Figure 1a). When the genetic code 4 block follows the standard genetic code block (blue), Mgcod correctly predicted the sequence of blocks in 98.4% of the cases with a mean shift of 47 bp and a standard deviation of 4634 bp. In 88.6% of the cases, Mgcod predicted the switch position with a shift of less than 5000 bp. When the standard genetic code block follows the genetic code 4 block (green), Mgcod predicted the correct sequence of changes in genetic codes in 98.0% of cases with a mean shift of -374 bp and a standard deviation of 5174 bp. In 85.4% of cases, Mgcod predicted the switch location with an error of less than 5000 bp. The negative sign of the mean error suggests that Mgcod is likely to predict the switch upstream to the true switch location.

**Figure 1:**
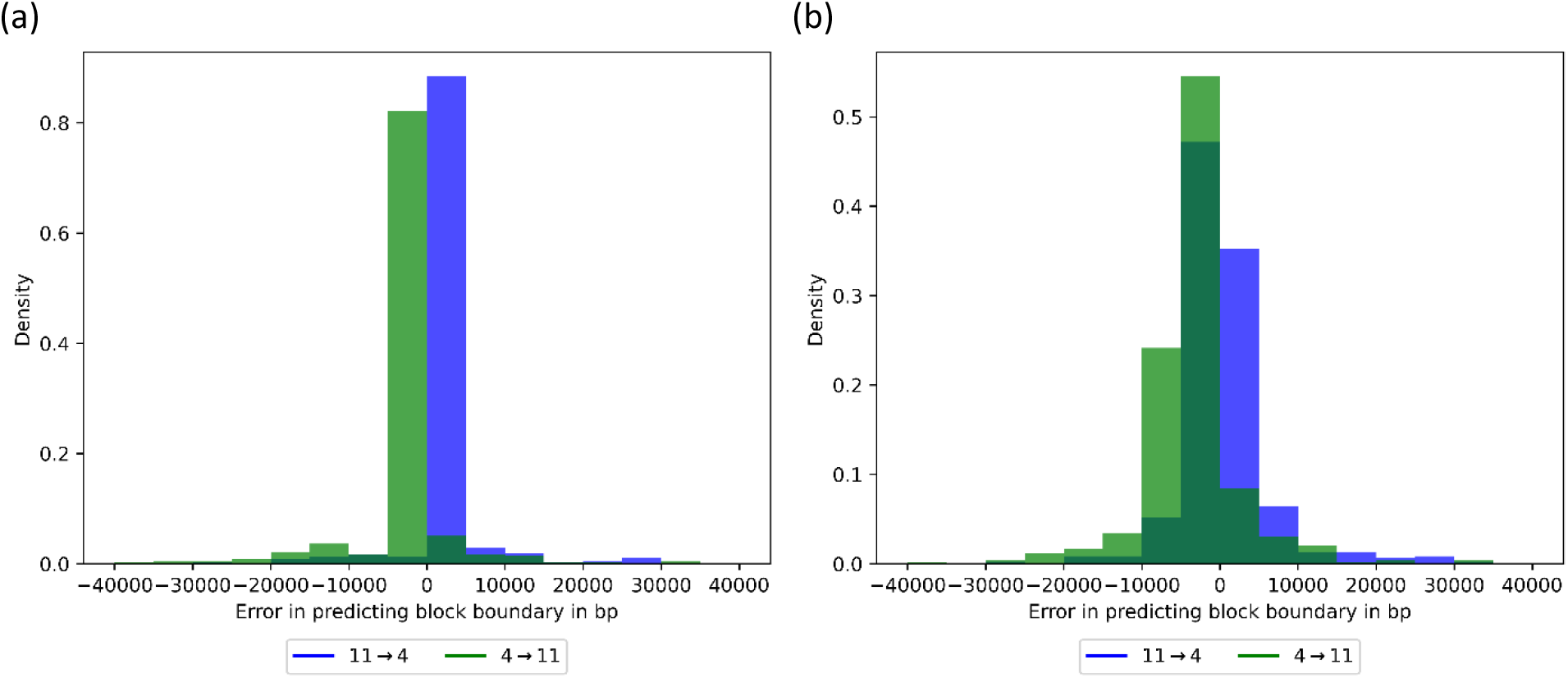
Histograms of the shifts in predicting the switch of genetic codes in simulated genomes with blocks with standard genetic code and with genetic code 4 **a)** with simulated change of protein-encoding strand and **b)** without simulated change of protein-encoding strand

To determine to which extent predictions of switch positions rely on PES information, we evaluated the performance of Mgcod without simulating a PES switch. As expected, the errors increased. The predictions had shifts from true positions of switches in the ranges -2170 ± 5631 bp when the block with standard genetic code was followed by the block with genetic code 4 and -867 ± 6251 bp when the blocks were situated in the opposite order (Figure 1b). Furthermore, the predicted block boundaries were within 5000 bp in 68.97% and 75.94% from the true positions, respectively, depending on the above-mentioned order of the blocks.

### Finding metagenomic contigs with multiple genetic codes

We screened 361,992 metagenomic contigs with a length between 80 kb and 300 kb for stop codon reassignments. We identified several types of pairs of genetic codes (Table 2). Since we were interested in contigs of phage origin, we applied several filtering steps (see Materials and Methods). In human (gut) metagenomes, we identified 45 phage contigs with standard genetic code (code 11) and genetic code 4, 694 contigs with code 11 and code 15, and 230 contigs with code 11 and code 101. Of those three groups of contigs, 17, 303, and 88 contigs, respectively, appeared to be complete based on the circularity feature, i.e., an exact match of 50 – 200 bp at the contigs ends (Table 2). More detailed information on the phage contigs with multiple genetic codes found in human (gut) metagenomes are given in Table S5. By using MSA, we found evidence supporting the predicted *opal* and *amber* stop codon reassignments in the likely complete phage genomes. However, we did not find an MSA support for the predicted *ochre* stop codon reassignments (Materials and Methods). In genomes with code 11 and code 4, in-frame *opal* stop codons most frequently aligned with themselves or UGG, which encodes tryptophan. In genomes with code 11 and code 15, in-frame *amber* stop codons most frequently aligned with themselves or a codon encoding glutamine. The observation that reassigned stop codons often align with themselves suggests that the reassignments have been conserved in evolution. In genomes with code 11 and code 101, in-frame *ochre* stop codons were most frequently aligned with themselves or with one of the two remaining stop codons, suggesting an incorrectly predicted recoding (Figure S1).

**Table 2:** The numbers of contigs with multiple genetic codes found in metagenomes.

|  | 11/4 | 11/15 | 11/101 | 4/15 | 4/101 | 15/101 |
| --- | --- | --- | --- | --- | --- | --- |
| **Number of contigs** | 3028 | 3985 | 9415 | 48 | 22 | 13 |
| **Contigs with $\leq 2$ genetic code switches** | 2887 | 3540 | 9269 | 36 | 16 | 7 |
| **Shorter block makes at least 20% of the contig** | 683 | 1519 | 1686 | 23 | 6 | 4 |
| **Human (gut)** | 288 | 840 | 1289 | - | 1 | 1 |
| **Viral** | 45 | 694 | 230 | - | - | - |
| **Circular** | 17 | 303 | 88 | - | - | - |

Previously, Yutin et al. identified 596 crAss-like phages among metagenomic contigs by searching circular contigs for the presence of phage marker genes. The authors reported *opal* or *amber* stop codon reassignments in 39 and 229 crAss-like phages, respectively. They also identified an additional phage with a not well-defined reassignment (Yutin et al. 2021). In this study, we rediscovered many of the crAss-like phages with stop codon reassignments identified by Yutin et al. (Table S5). Of the 17 likely complete phage genomes with code 11 and code 4, eleven were identified as crAss-like phages by Yutin et al., and all were predicted with genetic code 4. Of the 303 putatively complete phage genomes with code 11 and code 15, 211 were identified as crAss-like phages by Yutin et al. They predicted that 198 of these genomes employ genetic code 15 and identified that some kind of reassignment was present in yet another genome. For the remaining 12 crAss-like phages, in which we identified code 11 and code 15, Yutin et al. predicted genetic code 11. We determined that in these 12 genomes, the blocks of code 15 had genes that rendered highly similar protein sequences to translations of genes annotated by code 11. Of the 88 likely complete genomes with code 11 and code 101, Yutin et al. also identified 70 of those phages genomes as crAss-like but predicted all of them to use the standard genetic code. This was in line with our previous conclusion made from the MSA analysis that these predictions were likely false positives. Therefore, the genomes of phages with standard genetic code and genetic code 101 were excluded from subsequent analysis.

The significant overlap of likely completely assembled phage genomes with known crAss-like phages suggests that many of the incomplete contigs with multiple codes, code 11 and code 4 (or code 15), are also likely crAss-like phages. This statement is also supported by the observation that ∼86% (24/28) of the incomplete phage genomes with code 11 and code 4 had an average nucleotide identity (ANI) ≥ 0.8 to at least one likely complete crAss-like phage with the same type of multiple genetic codes. The average pairwise ANI of likely incomplete phage genomes with code 11 and code 4 to a sequence of the best matching complete crAss-like phage was 0.965 ± 0.012 (Table S6). Similarly, ∼90% (352/391) of the incomplete phage genomes with code 11 and code 15 had an ANI ≥ 0.8 to at least one likely complete phage genome, with an average ANI with the best matching complete crAss-like phage being 0.970 ± 0.033 (Table S7).

### Biological meaning of Mgcod predictions

To assess whether the segmentation predicted by Mgcod is biologically meaningful, we interrogated 320 genomes of likely complete phages predicted to have multiple codes, code 11 and code 4 (or code 15). Considering the patterns in functional annotations of protein-coding regions and the locations of predicted suppressor tRNA genes, we argue that the Mgcod-defined blocks are biologically meaningful.

We found that genes without stop codon reassignments encode early-stage phage genes, e.g., genes involved in DNA replication and transcription (Tables S8 and S9), while genes harboring reassigned in-frame stop codons are involved in late-stage processes such as virion assembly (Tables S10 and S11) (Ivanova et al. 2014; Yutin et al. 2021; Borges et al. 2021).

While only three out of the 17 likely complete phage genomes with code 11 and code 4 encode suppressor tRNA with an anticodon corresponding to the *opal* stop codon (Sup-TCA tRNA), 225 out of the 303 likely complete phage genomes with code 11 and code 15 carry a suppressor tRNA with an anticodon corresponding to the *amber* stop codon (Sup-CTA tRNA). Furthermore, two of the three phage genomes with code 11 and code 4 that encode Sup-TCA tRNA are also encoding a Sup-CTA tRNA, and eight genomes with code 11 and code 15 encode two Sup-CTA tRNAs (Table S12). The absence of suppressor tRNAs in the remaining genomes may indicate that the phages rely on a host-encoded suppressor tRNA, encode a non-canonical tRNA not predicted by current methods, or simply that the genomes are incomplete.

In phages with alternating code 11 and code 15, the Sup-CTA tRNA gene is usually located at the 5’ end of the PES that carries genes with *amber* stop codon reassignment (Figure 2).

**Figure 2:**
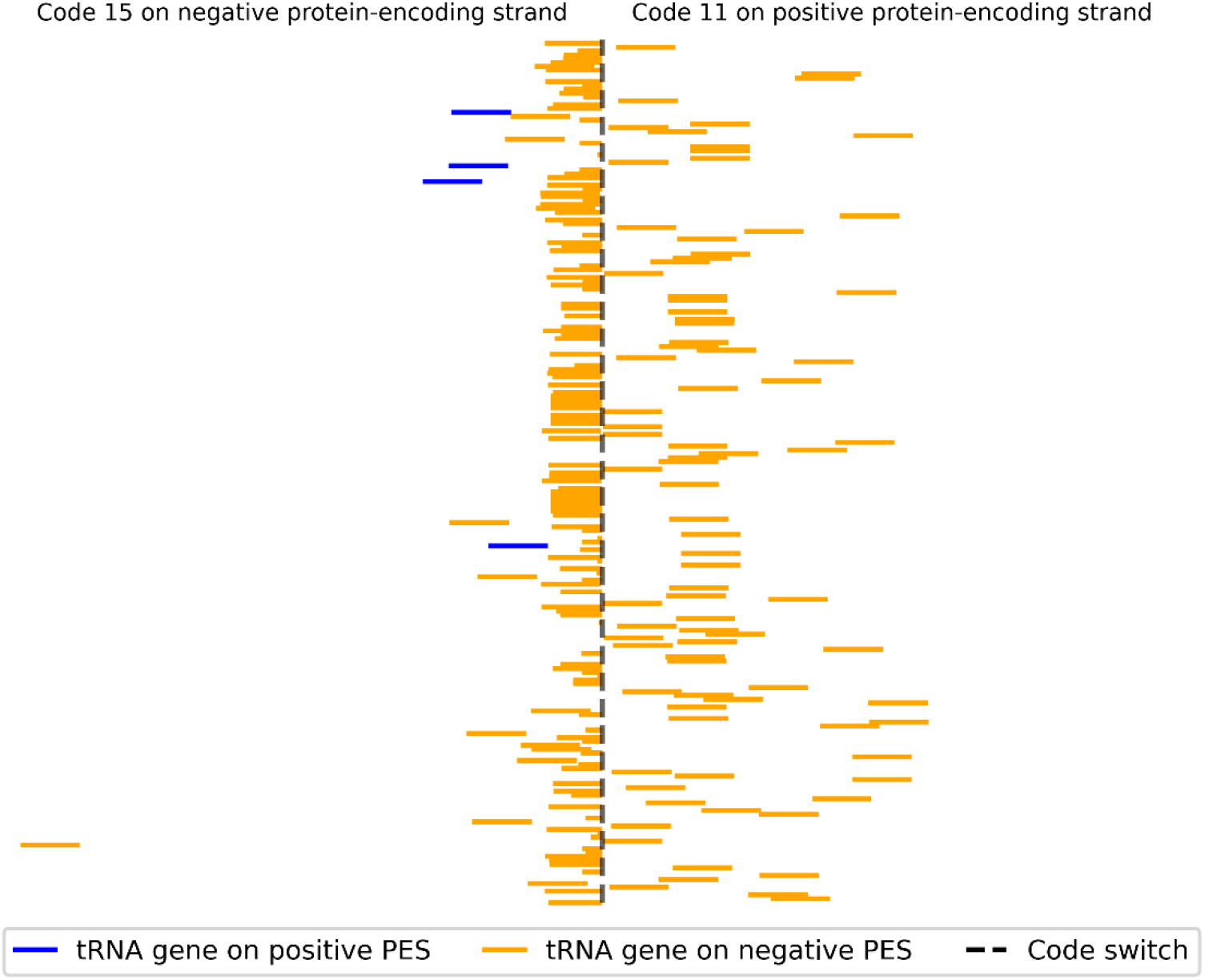
Distribution of locations of Sup-CTA-tRNA genes with respect to the predicted switch point of genetic codes in 303 likely complete phage genomes with blocks of code 11 and code 15. The Sup-CTA-tRNA is usually located at the 5’ end of the protein-coding strand carrying genes encoded in genetic code 15. Possibly, the predicted switch point was slightly misplaced when Sup-CTA-tRNA was predicted in the negative protein-encoding strand but falls into a genomic region predicted to encode genes in code 11. In this figure, the lengths of protein-coding strands were normalized to 1.

Assuming sequential expression of the genes, this configuration implies that the Sup-CTA-tRNA is expressed before the genes with stop codon reassignments, ultimately enabling their expression. Thus, the set of functions of predicted protein-coding genes and the location of suppressor tRNA show that the genome segmentation of Mgcod is biologically meaningful.

### One block in phages with multiple genetic codes has dual-coding

In our analysis of likely complete phage genomes, we found that one block is usually dual-coded, i.e., two models utilizing different genetic codes produce almost equivalent proteins in this block (Table S5). In phage genomes with blocks of code 11 and code 15, the coding regions in the block with code 11 are usually also predicted by the model using code 15. Conversely, only in a few cases, the coding regions in the block predicted to employ genetic code 15 are dual-coded (Figure S2). In phage genomes with code 11 and code 4, blocks with code 11 and code 4 are dual-coded with equal frequency. However, the analysis is confounded by the small number of genomes with alternating code 11 and code 4 (Figure S2).

To give an example, we show results of the analysis of Phage 2 that alternatively employs code 11 and code 15 (Fig. 3). Mgcod’s segmentation of Phage 2 into two blocks is consistent with the segmentation done by Ivanova et al. (2014) using the same genome linearization point (see Fig. 4 in Ivanova et al. (2014)). However, we determined that while genes in one block were distinctly translated by genetic code 15 (red), the other block was dual-coded (grey). Genes predicted only by the model with code 11 (blue) are present in both blocks. Most of these predictions are short and have low log-odds scores, which is typical for false-positive predictions. We did not filter out these predictions as false positives could be easily excluded in downstream analyses.

**Figure 3:**
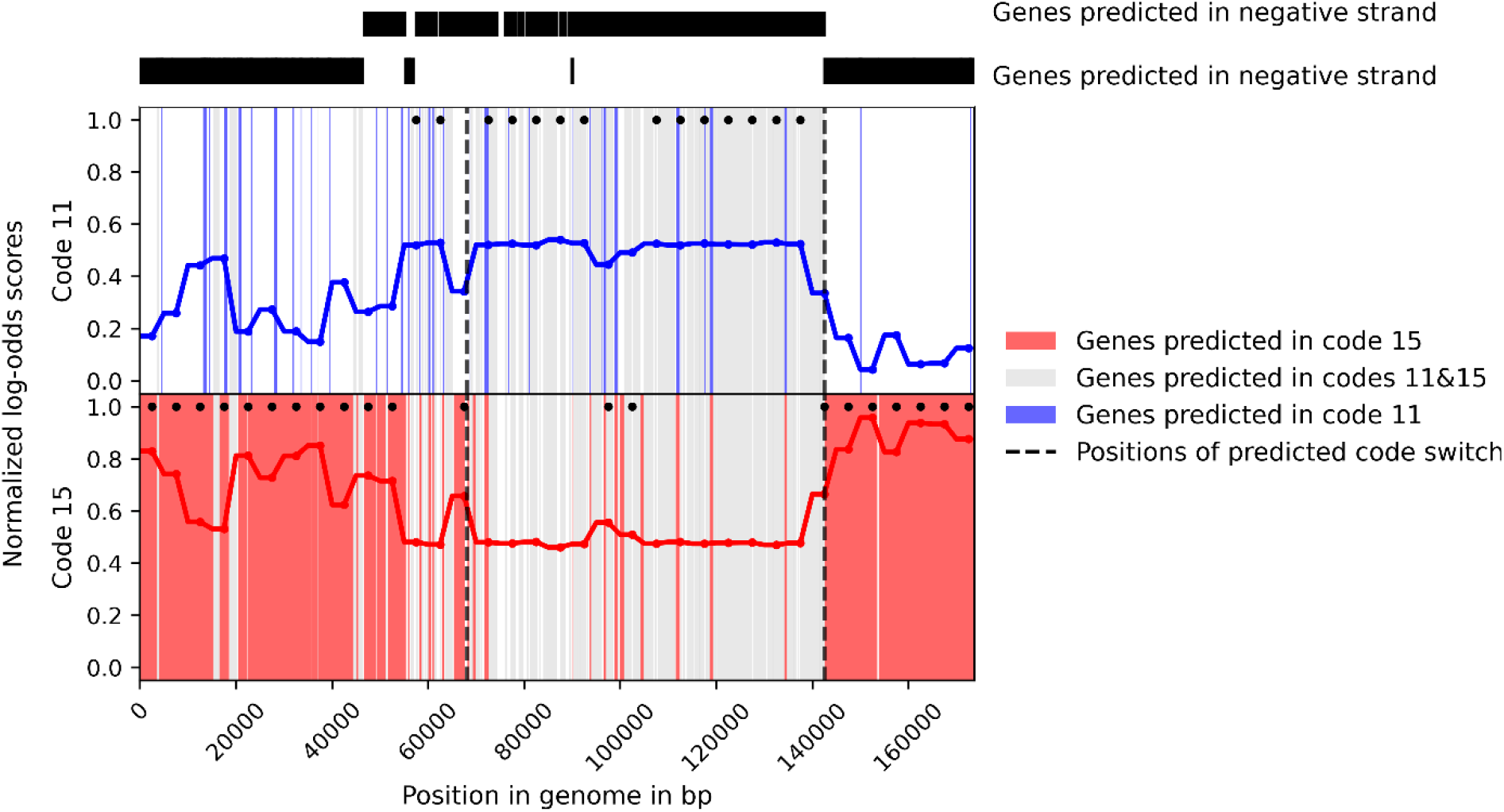
Visualization of change of predicted genetic code along the genome of Phage 2 (Ivanova et al. 2014). The top of the Figure shows the DNA strands of predicted coding regions in black. The colored curves in the two panels below show the normalized values of the genes log-odds scores computed by the two MGM models in 5 kb long windows. The type of a model with a higher score is indicated by a dot at the top of the corresponding panel. Blocks boundaries are indicated by the vertical dashed lines. Two blocks of genes encoded with code 15 are seen in the negative strand. (Since the genome is circular, these two blocks are parts of a large continuous block split by the genome linearization point. Another block (in the middle) is dual-coded, i.e., translations of predicted coding regions made with either genetic code are almost identical (grey color).

Furthermore, functional annotations of protein-coding regions that could be translated by either code with little difference at a protein level show that dual-coded genes are enriched for phage genes involved in early-stage DNA replication and transcription (Tables S13 and S14). Therefore, these genes can be translated using the machinery of a host with the standard genetic code while a switch of the genetic code does not disrupt their translation.

## Discussion

Here, we presented Mgcod, a tool that identifies a type of the genetic code in a metagenomic contig – even if more than one genetic code is present - and provides annotation of protein-coding regions. Independent identification of the genetic code in parts of the contig enables more accurate annotation of coding regions.

In simulated genomes with multiple genetic codes, Mgcod accurately segmented genomes into genomic blocks. However, the simulated genomes are idealized subjects with two distinct genetic codes. Our search through metagenomic databases for phage contigs with multiple genetic codes demonstrated that the reality is more complex. The distinction of blocks with different genetic codes is not always as clear as in simulated genomes since one block is usually dual-coded, i.e., coding regions can be translated with either genetic code yielding nearly identical proteins. This dual coding makes the problem more difficult. In some instances, the genes and proteins are so similar that it is impossible to determine the true genetic code *ab initio*. Therefore, this type of genome organization may lead to false-positive predictions of multiple genetic codes, e.g., alternating code 11 and code 101. Despite the complication due to the dual coding, annotation of protein-coding regions and the localization of corresponding suppressor tRNAs genes showed that Mgcod predicts block structure of phage genomes with multiple genetic codes in biologically meaningful ways.

Mgcod appears to be a useful tool for screening metagenomes and discovering novel phages with multiple genetic codes. While we initially identified contigs with several types of multiple genetic codes, after additional filtering, only contigs with code 11 and code 4 (or code 15) were retained. We have identified 34 and 483 previously unknown crAss-like phages in human (gut) metagenomes with code 11 and code 4 (or code 15).

Prediction of dual coding blocks in phage genomes has implications for hypothesized infection mechanisms of such phages. For instance, crAss-like phages with multiple genetic codes are predicted to infect microbial species of phylum *Bacteroidetes*, which use the standard genetic code (Dutilh et al. 2014; Guerin et al. 2018; Yutin et al. 2018; Edwards et al. 2019; Koonin and Yutin 2020; Yutin et al. 2021; Benler et al. 2021). One proposed mechanism of infection builds on the observed PES bias and the absence of stop codon reassignments in early-stage phage genes. According to this mechanism, early-stage genes are translated with the standard genetic code using the host’s translation machinery, producing proteins, e.g., release factors, which enable the switch of genetic code later in the infection cycle (Ivanova et al. 2014). Another proposed mechanism is that the phages use bacterial tRNA synthetases to charge the suppressor tRNA, and its presence is sufficient to enact a switch in genetic code (Borges et al. 2021). In favor of the later mechanism speaks the localization of the suppressor tRNA gene at the 5’ end of the PES containing late-stage phage genes with stop codon reassignment. Both hypotheses imply that a switch of genetic codes disrupts the translation of early-stage phage genes (Ivanova et al. 2014; Yutin et al. 2021; Borges et al. 2021). However, there might not be such a “hard” switch of genetic code as early-stage phage genes are usually dual-coded. Instead, our results suggest that there might be a “soft” switch, enabling the translation of recoded late-stage phage genes without disrupting the translation of dual-coded early-stage phage genes. Given the presence of differences in translations of the dual-coded genes, we provide annotations of such genes using the concept of isoforms. We caution, though, that wet-lab experiments are necessary to prove that both isoforms are expressed *in situ*.

Despite being designed to annotate contigs with presumably multiple genetic codes, Mgcod can also annotate any prokaryotic sequence with or without stop codon reassignments. Thus, Mgcod can be used as a tool for the annotation of coding regions in metagenomic sequences. The significance of such a tool is increasing as more contigs with stop codon reassignments are discovered in the course of ongoing sequencing. Obviously, off-the-shelf gene annotation tools may not render accurate gene predictions in contigs with non-standard and especially multiple genetic codes. In contrast, Mgcod starts with machine learning identification of the genetic code and applies the predicted codes, either block specific or not, to annotate protein-coding regions.

## Acknowledgements

We thank Tomas Bruna and Karl Gemayel for their helpful discussions. The work was supported by a scholarship from the German American Fulbright Commission to A.P. and the NIH grant GM128145 to M.B.

