## supplementary figures and table legends for "Annotation of Phage Genomes with Multiple Genetic Codes"

**Algorithm S1:** Recognition of genetic codes and alternating genetic codes in phage genomes and annotation coding regions.

---

**Input:** FASTA file
**Output:** GTF/GFF2 formatted gene annotations that account for the genetic code

```
 1. Run MetaGeneMark with models of protein-coding region with genetic code 11,
    4, 15, and 101
 2. If not predict alternating genetic codes
       a. Assign genetic code to input sequences based on model of protein-coding
          regions with highest coding potential and provide corresponding gene
          annotation
    Else
       a. Apply sliding window approach (5000 bp)
           i. Determine model with highest coding potential (log-odds scores) in
              each window
          ii. Assign window labels accordingly
       b. If a model has the highest potential in all windows
           i. This model's predictions are used as gene annotations in the whole
              genome
         Else
           i. Identify the two models that have most frequently the highest
              coding potential in a window (A & B)
          ii. Segment genome into genomic blocks
               1. Set longest consecutive sequence of windows, which have the same
                  window label and have not been updated, as seed block;
                  Set corresponding label as reference
               2. Extend seed block into both direction until n consecutive
                  mismatches are observed
               3. Update labels of extended sequence of windows according to
                  reference
               4. Repeat a-c until all window labels have been updated
               5. Define tentative block boundaries as the center of intergenic
                  region separating the two blocks
               6. If the PES switches within one window of predicted block boundary,
                  update prediction to center of intergenic region separating the
                  genes between which the PES switches
         iii. Compile set of predicted protein coding regions
               1. For all predictions made by models A & B
                   a. If model A predicts a long gene and model B many short genes
                      on the same PES, and coding potential of model A ≥ model B
                       1. Keep prediction of model A; Drop predictions of model B;
                          Assign gene label A
                   b. Elif model A predicts a long gene and model B many short genes
                      on the same PES, and coding potential of model A < model B
                       i. Drop prediction of model A; Keep predictions of model B
                          Assign gene label B
                   c. Elif predictions of model A & B are identical
                       i. Keep either prediction
                          Assign gene label C
                   d. Elif (predictions of model A and B share stop coordinates) or
                      (share start coordinates and stop coordinates differ by less
                      than t bp)
                       i. Keep both predictions and annotate as isoforms
                          Assign gene label C
                   e. Else
                       i. Keep prediction
                          Assign gene label A or B accordingly
```

---

### Supplementary Figures

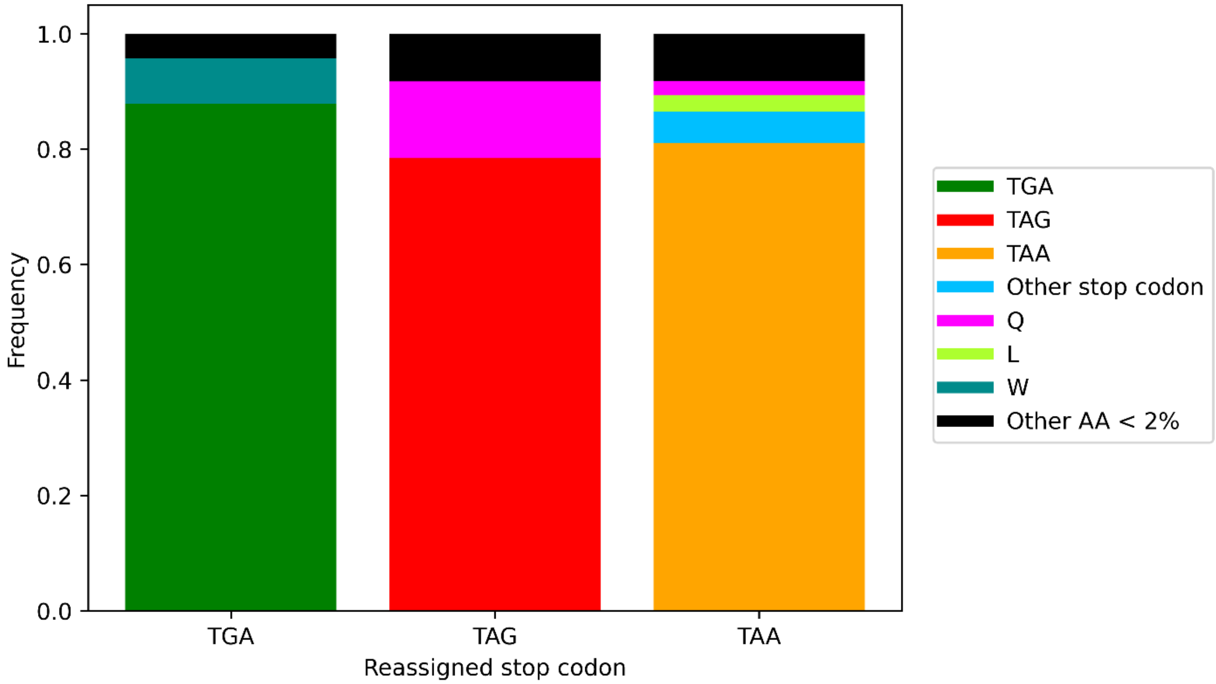


**Figure S1:** Frequencies with which reassigned stop codons aligned to codons for particular amino acids (or the same likely reassigned stop codons) in MSA of homologous nucleotide sequences. The alignment evidence supported predicted reassignments of the *opal* stop codon (TGA) in phages with alternating codes 11 and 4. Similarly, there was supporting evidence for the predicted reassignment of the *amber* stop codon (TAG) in phages with alternating codes 11 and 15. Since the TAA stop codon most frequently aligned with other stop codons and rarely to sense codons, the predicted reassignments of the *ochre* stop codon (TAA) in phages with alternating code 11 and code 101 were likely false.

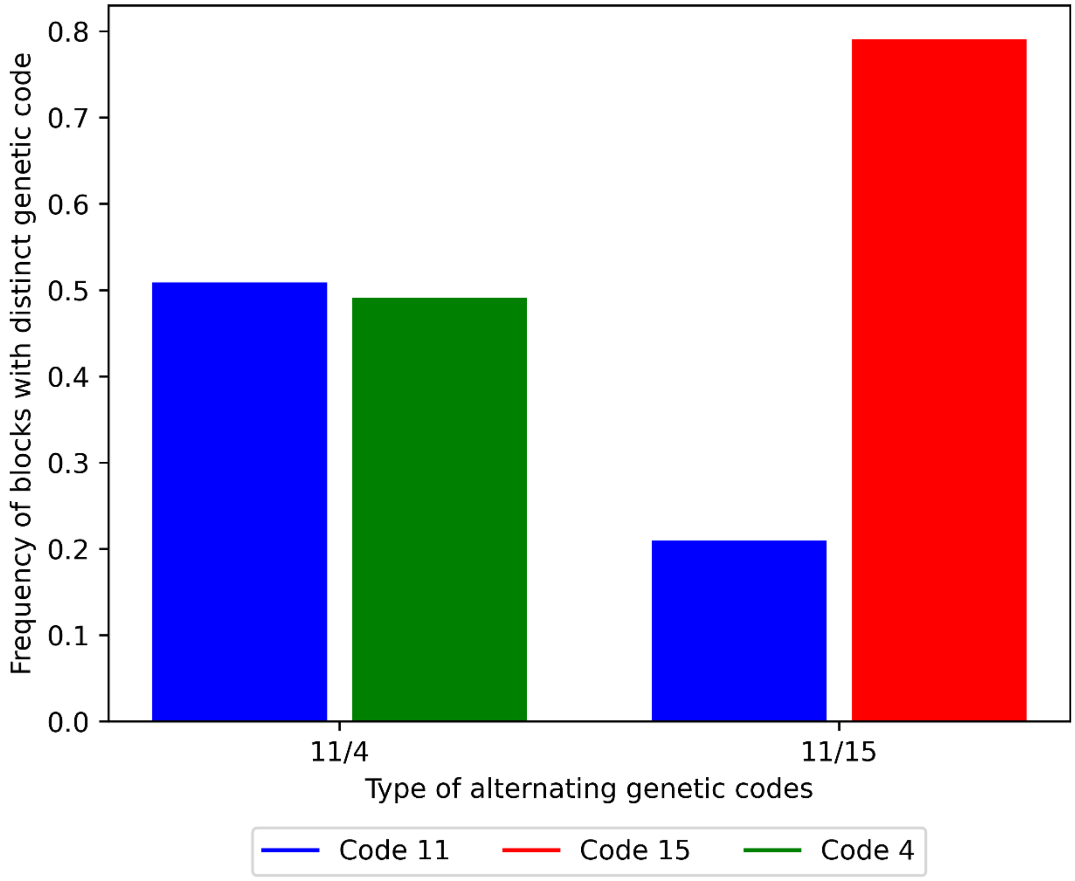


**Figure S2:** Frequency of types of blocks predicted in phage genomes which contained a block with distinct genetic code and a dual-coding block. In the 45 phage genomes with alternating code 11 and code 4 (11/4), the block employing a distinct genetic code was equally frequently encoded with code 11 and code 4. In contrast, in the 694 phage genomes with alternating code 11 and code 15 (11/15), the block employing a distinct genetic code was more frequently encoded with code 15.

### Supplementary Table Legends

**Table S1:** A list of used representative bacterial genomes with genetic code 4/25.

**Table S2:** A list of used representative bacterial genomes with genetic code 11 with a size smaller than 2Mb and GC% less than 35%.

**Table S3:** A list of used representative bacterial genomes with genetic code 11 and with a genome size greater than 2Mb and GC% greater than 35%.

**Table S4:** Accuracy of the GCD concerning the identification of genetic code in contigs of various lengths. *Opal* stop codon reassignments in genomes greater than 2Mb and GC% greater than 35% as well as *amber* and *ochre* stop codon reassignments were simulated. The genomes were then split into contigs of various lengths, and the accuracy was evaluated on 1000 randomly selected contigs.

**Table S5:** Summary of viral contigs with multiple genetic codes found in human (gut) metagenomes. Completeness was assessed based on circularity, i.e., exact of 50 - 200 bp between the ends. If the contig was also identified as crAss-like by Yutin et al. (2021), their genetic code prediction is also reported.

**Table S6:** Contigs with blocks of code 11 and code 4 that were classified as incomplete (column A) have a high average nucleotide identity with a complete phage with blocks of code 11 and code 4 (column B). The pairwise average nucleotide identity (ANI) is shown in column C. Only matches with ANIs ≥ 0.8 are reported.

**Table S7:** Contigs with blocks of code 11 and code 15 that were classified as incomplete (column A) have a high average nucleotide identity with a complete phage with blocks of code 11 and code 15 (column B). The pairwise average nucleotide identity (ANI) is shown in column C. Only matches with ANIs ≥ 0.8 are reported.

**Table S8:** Functional annotation of cluster centroids of coding regions annotated with code 11 in phages with code 11 and code 4. Cluster centroids were functionally annotated based on homology to clusters of orthologous groups included in the prokaryotic Virus Orthologous Groups (pVOGs) database.

**Table S9:** The same as in Table S8 for cluster centroids of coding regions annotated with code 11 in phages with code 11 and code 15.

**Table S10:** The same as in Table S8 for cluster centroids of coding regions annotated with code 4 in phages with code 11 and code 4.

**Table S11:** The same as in Table S8 for cluster centroids of coding regions annotated with code 15 in phages with code 11 and code 15.

**Table S12:** Predicted suppressor tRNAs in putative complete phage genomes with blocks of standard genetic code and genetic code 4 (or code 15).

**Table S13:** Functional annotation of cluster centroids of dual-coded coding regions in phages with genetic code 11 and code 4.

**Table S14:** Functional annotation of cluster centroids of dual-coded coding regions in phages with genetic code 11 and code 15.
